# Dynamics of positional information in the vertebrate neural tube

**DOI:** 10.1101/2024.07.01.601514

**Authors:** Anđela Marković, James Briscoe, Karen M Page

## Abstract

In developing embryos, cells acquire distinct identities depending on their position in a tissue. Secreted signaling molecules, known as morphogens, act as long-range cues to provide the spatial information that controls these cell fate decisions. In several tissues, both the level and the duration of morphogen signaling appear to be important for determining cell fates. This is the case in the forming vertebrate nervous system where antiparallel morphogen gradients pattern the dorsal-ventral axis by partitioning the tissue into sharply delineated domains of molecularly distinct neural progenitors. How information in the gradients is decoded to generate precisely positioned boundaries of gene expression remains an open question. Here, we adopt tools from information theory to quantify the positional information in the neural tube and investigate how temporal changes in signaling could influence positional precision. The results reveal that the use of signaling dynamics, as well as the signaling level, substantially increases the precision possible for the estimation of position from morphogen gradients. This analysis links the dynamics of opposing morphogen gradients with precise pattern formation and provides an explanation for why time is used to impart positional information.

## Introduction

The process of tissue development is remarkably reliable and precise. It involves the generation of distinct cell types in a characteristic and reproducible arrangement to form organised patterns of cell fates. In an influential essay that describes this process, Lewis Wolpert introduced the concept of “positional information” (1). This proposed that positional information is derived from signals, usually referred to as morphogens, which form concentration gradients in the tissue, with cells measuring the local concentration and interpreting it to select the appropriate cell fate for that position. In this view, concentration thresholds define boundaries that distinguish adjacent sets of cell types.

The dorsal-ventral patterning of the vertebrate nervous system is a well established system for studying morphogen-driven patterning (2). Signals emanating from two opposing sources – the dorsal roof plate secreting the morphogen BMP, and the ventral floor plate secreting the morphogen Shh – form antiparallel gradients that partition the neural tube into 11 discrete domains of molecularly distinct neural progenitors arrayed along the dorsal-ventral axis, Fig. 1. Neural progenitors respond to morphogen signaling by controlling a gene regulatory network that specifies the pattern of gene expression along the dorsoventral axis and hence the positions at which distinct neuronal subtypes are generated. Gene expression and consequently cell fate appear to be specified by a combination of Shh and BMP signaling (3). Positions of several gene expression boundaries have been experimentally measured (3–6) and this indicates that the boundaries are specified with a high degree of precision, resulting in sharply defined transitions in gene expression and only moderate intermixing of cell types at domain boundaries.

**Fig. 1.**
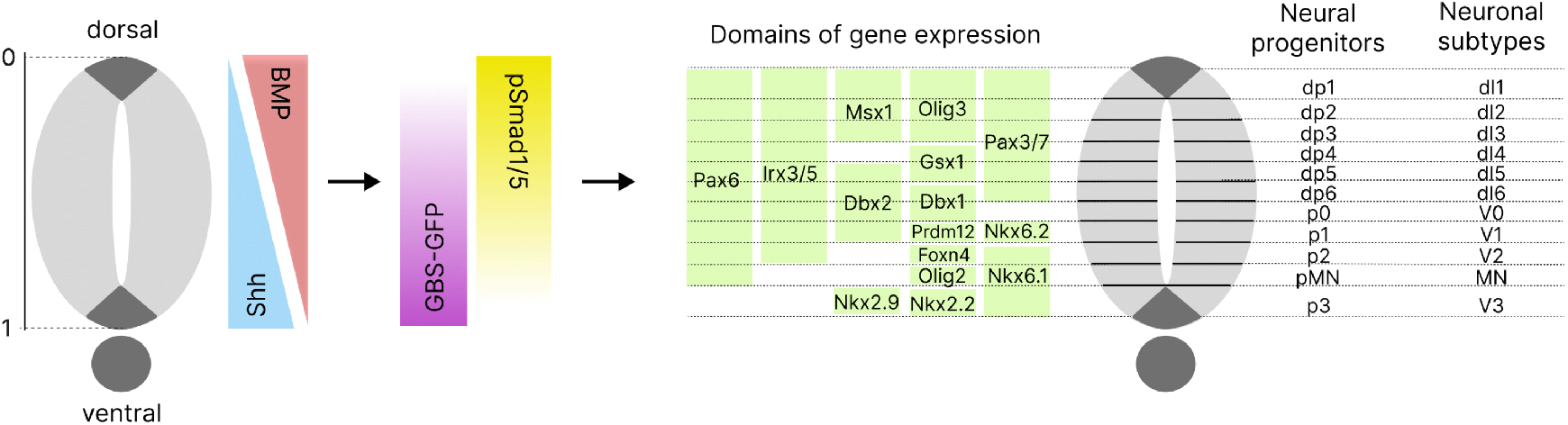
The morphogen, Shh, emanating from the ventral pole, and the morphogen, BMP, secreted from the dorsal pole of the neural tube, form antiparallel gradients along the dorsal-ventral axis. These extracellular gradients are translated into intracellular gradients of the activity of Gli and Smad transcription factors, regulated by Shh and BMP signaling, respectively. Their activity is measured using GBS-GFP, a Gli reporter, and pSmad1 / 5, the activated version of Smad1/5. The Gli and Smad transcription factors control gene regulatory networks that specify the domains of gene expression along the axis, and hence, partition the neural tube into 11 discrete domains of molecularly distinct neural progenitors. In this way, the positions at which distinct neuronal subtypes will be generated are determined.

To explore the amount of positional information that can be obtained from the two morphogens, we took advantage of tools from *information theory* (7). These have been used previously to study information transmission through signaling pathways and gene regulatory networks (8–16), including positional information transmission by the morphogen Bicoid, which governs the anterior-posterior patterning of the Drosophila blastoderm (17–21). The variability between embryos of the Bicoid gradient and the expression levels of the genes it controls was used to connect positional information to experimental data at a specific developmental time point. This revealed that Bicoid has a lower precision than its target genes (17), leading to the possibility that other sources of information or mechanisms are involved, such as continuous, rather than instantaneous assessment of Bicoid concentration (22).

The Drosophila blastoderm differs in several ways from many developing tissues. It is a syncytium. This allows Bicoid to regulate gene expression directly without an intervening signal transduction pathway. Most developing tissues, including the neural tube, are cellularised and morphogens spread extracellularly to activate intracellular signaling pathways that control gene expression (23, 24). In addition, the patterning of the blastoderm occurs in the absence of tissue growth (25). Most tissues grow as they are patterned, and morphogen gradients do not always scale with growth (6, 23). This, along with other factors such as changes in the production of the signal, results in the gradient changing as the tissue is patterned (23). Consequently, there is no straightforward correlation between the concentration of morphogens, their location in tissue, and cell fate allocation. Indeed, the duration of morphogen signaling has also been shown to influence cell fate decisions (4, 26–28). It is unclear, given the dynamics and the heterogeneity and fluctuations in the spread and interpretation of a morphogen, how a graded morphogen produces the precise patterns of cell fate decisions that are observed in developing tissues.

Despite the prominence of Wolpert’s positional information concept, it has become evident that relying solely on morphogen concentration does not provide a comprehensive explanation for understanding morphogen gradients and their role in tissue development (24). The “progressive and dynamic manner” (26) in which the domains of neural progenitors appear suggests new ways of encoding positional information. Here, we set out to extend the information-theoretic approach by adapting methods that accommodate dynamics to analyse positional information in the neural tube. We wanted to explore the impact of signaling dynamics on positional precision and introduce a temporal dimension to the concept of positional information.

## Results

### Quantifying positional information with information theory

We first set out to quantify how much positional information the Shh and BMP morphogens provide in the neural tube. The signaling pathways of these morphogens represent complex, noisy channels through which the cells receive information about their positions (23). Information theory describes the input and the output of any noisy channel as two random variables, *X* and *Y*, respectively (29). The noise in the channel leads to a distribution of possible output responses produced by the same input message (16), and the uncertainty in the input arises from the noise in its source as well as other features of the system. Let *X* be a discrete random variable that can take values *x*_1_, *x*_2_, …, *x*_*m*_, with the probability distribution *P*_*X*_ (*X*) = (*P* (*x*_1_), …, *P* (*x*_*m*_)), and let *Y* be a continuous random variable, with the probability distribution *P*_*Y*_ (*Y*) and 𝒴 as a space of all possible values of *Y*. The main measure of information that we use here, *mutual information* between two random variables *X* and *Y, MI*(*X, Y*), is defined by (30)

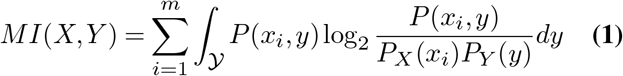

where *P* (*x*_*i*_, *y*) = *P* (*y* |*x*_*i*_)*P*_*X*_ (*x*_*i*_) = *P* (*x*_*i*_| *y*)*P*_*Y*_ (*y*). Mutual information *MI*(*X, Y*) measures to what extent *X* and *Y* depend on each other, being 0 when they are independent, and captures any kind of correlation between the input and the output (19). It indicates how much the uncertainty about the value of the random variable *X* is reduced by knowing the state of *Y*, that is, how much information about the input we obtain after looking at the output (29). More precisely, *MI*(*X, Y*) can be interpreted as the average number of binary (yes/no) questions about the value of *X* that are answered after observing *Y*. Each question divides possible values of *X* into two equally probable, mutually exclusive classes based on the values of *Y* associated with them.

Hence, 2^*MI*(*X,Y*)^ gives us the average number of equiprobable classes of the values of the input *X* that produce classes of the values of the output *Y* that can be distinguished without error (31). Therefore, the maximum value of mutual information in this case is log_2_ *m*, when all input values can be perfectly resolved. Two individual values of *X, x*_*i*_ and *x*_*j*_, are said to be resolved if their corresponding output distributions *P* (*Y* |*X* = *x*_*i*_) and *P* (*Y*| *X* = *x*_*j*_) do not overlap (32). When all the input values have the same probability, *P* (*x*_1_) = … = *P* (*x*_*m*_), then the mutual information depends solely on the overlap between their output distributions and provides a measure of the noise which leads to these overlaps and prevents the system from perfectly resolving each input value.

The outputs of the Shh and BMP signaling pathways are the activities of their intracellular transcriptional effectors. These were measured in the mouse neural tube by Zagorski et al. (3). Shh signaling results in the intracellular activation of Gli transcription factors, the activity of these was measured using a transcriptional reporter GBS-GFP. BMP signaling activates Smad transcription factors, which were measured using an antibody specific for phosphorylated SMAD1/5, pSmad1/5, the activated version of the BMP transcriptional effectors. Hence, we take the activities of GBS-GFP and pSmad1/5 to be the outputs of the Shh and BMP signaling pathways, respectively (Fig. 1). To quantify the positional information provided by the signaling of Shh and BMP morphogens in the neural tube, we computed *MI*(*X, Y*) where the input *X* represents different relative positions *x*_1_, …, *x*_*m*_ along the dorsal-ventral length of the neural tube, and the output *Y* the levels of GBS-GFP and pSmad1/5 activity measured at these positions (Fig. 2). Following the ideas in (17, 19), we argue that this definition of *MI*(*X, Y*) is a proper measure of positional information in the neural tube.

**Fig. 2.**
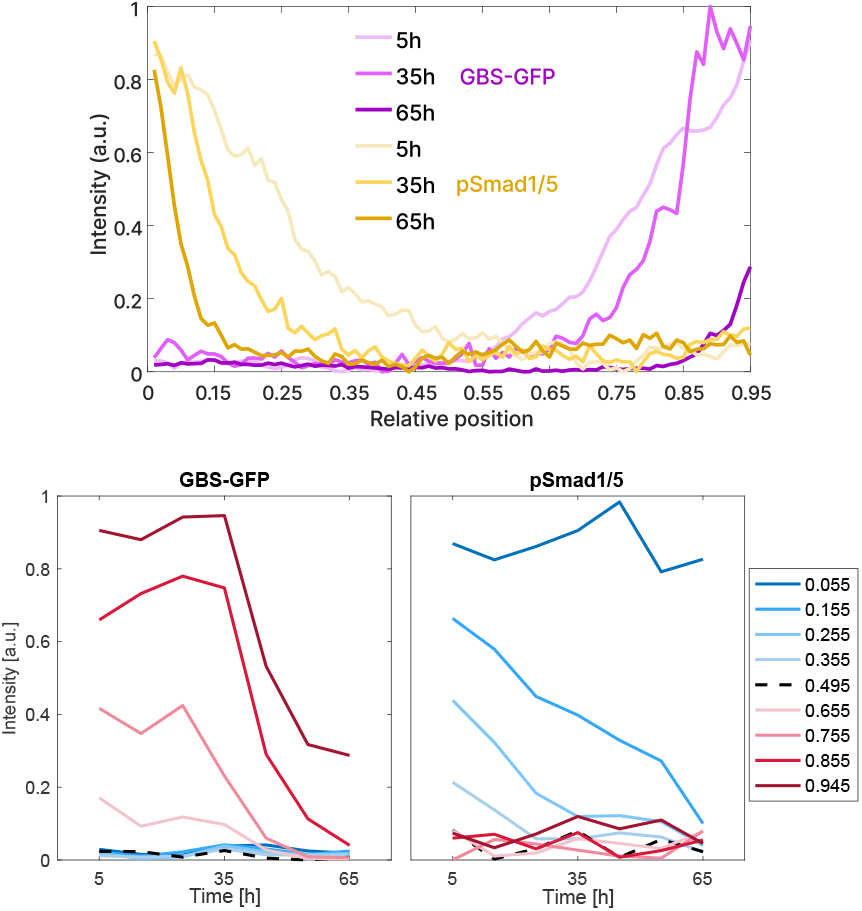
Top: the mean gradients of the activity of GBS-GFP, the reporter of Shh signaling, and the gradients of the activity of pSmad1/5, the transcriptional effectors of the morphogen BMP, along the relative dorsal-ventral length of the neural tube at 5h, 35h and 65h during the development. Bottom: the mean levels of GBS-GFP and pSmad1/5 over time at the indicated relative positions.

More precisely, the input probability distribution *P* (*X*) represents the positions of cells along the dorsal-ventral axis. Since cell density is approximately constant along this axis, we take the distribution *P* (*X*) to be uniform (6). The relative dorsal-ventral length of the neural tube from position 0.055 to 0.945, where the position 0 corresponds to the dorsal end and 1 to the ventral end, is divided into bins of width 0.05 with a distance of 0.01 between them. This resulted in 15 bins and therefore 15 input values denoted sequentially as *x*_1_, …, *x*_15_ (*Supplementary Information* (SI) Fig. 1). Using the measurements of the neural tube length over time from (3), the average cell diameter, 4.9*µ*m (6), is 0.05 of the relative length at 5h of neural tube development, the first time point at which the signals are measured. The absolute length of each bin grows over time, but during the first ∼30 hours of development, this length is less than 2 average cell diameters, and at 35h it is approximately 2 cells.

Zagorski et al. (3) measured the levels of GBS-GFP and pSmad1/5 every 0.01 of relative length from position 0.055 to 0.945, along the dorsal-ventral length of the neural tube in a single embryo. Hence, one of our 15 bins, *x*_*i*_, comprises 6 single relative positions at which the signals are measured and the output values corresponding to *x*_*i*_ are concatenated measurements of the signals made at those 6 positions in all embryos. As in Zagorski et al. (3) and Dubuis et al. (17), we focus on how much information there is at fixed relative positions in the tissue. The 15 bins allow us to determine this (see *SI* for details) and are independent of the progenitor domains, which are not of equal proportions and change during development (6).

Information-theoretic tools have been used before to study dynamic signals transmitting information to cells about their environment (33–35). This has led to the idea that a single cell monitors signals over time to overcome noise (33) and that information can be encoded in different dynamical patterns of signals (36). Therefore, the output values were represented as vectors of cellular responses at different time points. To explore the impact of morphogen signaling dynamics on positional precision, we took a similar approach and used the levels of GBS-GFP and pSmad1/5 measured at a particular relative position at successive time points, *t*_1_, *t*_2_, …, during neural tube development. Hence, a single output response to an input *x*_*i*_ becomes a vector of the form 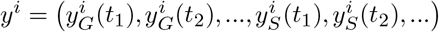 where G denotes GBS-GFP and S pSmad1/5. Note that the signals are only measured once in an embryo and then the temporal trajectory is inferred from measurements at equivalent positions in embryos of sequential time classes (see *SI* and (3, 6) for details).

To compute mutual information, we took advantage of a recently developed framework proposed by Jetka et al. (32) – SLEMI (statistical learning estimation of mutual information) – which simplifies the calculation of mutual information for high-dimensional output. The method involves a classifier that uses a logistic regression model to estimate *P* (*x*_*i*_| *y*), the probability of being at a position *x*_*i*_ after observing a signal level *y*, from the given data such that the probability of being at the true position for a given signal is maximised. Therefore, SLEMI is similar to “decoding-based information estimators”, described in (37), which use machine learning classifiers. We compared SLEMI with one of them, the decoding-based method proposed by Granados et al. (34). Jetka et al. (32) demonstrated the validity of their method and showed its advantages over the most commonly used method for computing mutual information, *k*-nearest neighbour (*k*NN). We also applied the *k*NN method to our data. The three computational tools give similar results (SI Fig. 2) and more detailed comparison can be found in *SI*. Overall, we believe that SLEMI is the right choice for our analysis.

The calculation of mutual information led to the values of 2^*MI*(*X,Y*)^ which, in this case, represent the average number of spatial regions along the dorsal-ventral axis of the neural tube that can be clearly delineated based on received signal levels. Since the positions are uniformly distributed, these regions are of equal length.

### Decoding Morphogen Gradients at Single Time Points

We computed mutual information (*MI*) between the positions along the dorsal-ventral axis of the neural tube and the levels of GBS-GFP and pSmad1/5 activity measured at these positions. We considered the levels of each signal separately and the combined levels. The data consist of 102 measurements of each of the two signals measured in each of the 15 spatial bins at each of the 7 time points 5h, 15h, 25h, 35h, 45h, 55h and 65h during neural tube development. We first calculated the mutual information for each time point by using only the signal levels measured at the corresponding time point. We then considered series of signal levels at successive time points and computed the time-series mutual information from the first time point up to each later time point. For example, the values measured at 5h, 15h, 25h and 35h were used to compute the mutual information for the 35h time course.

The values of time-point mutual information are highest at the first three time points, 5h, 15h, 25h, Fig. 3. However, when taking GBS-GFP and pSmad1/5 separately, the timepoint *MI* does not reach 1 bit, the value needed to distinguish two regions. This indicates that at a single point in time, on average, the information from either Shh or BMP signaling is insufficient to create a sharp boundary in the neural tube. On the other hand, the mutual information for the two signals combined is higher and shows that, at these first three time points, there is almost enough information to be able to resolve 3 regions (2^1.58^ ≈3). Thus, by sensing both morphogens at a single time point, there is sufficient information to specify 2 spatial regions.

**Fig. 3.**
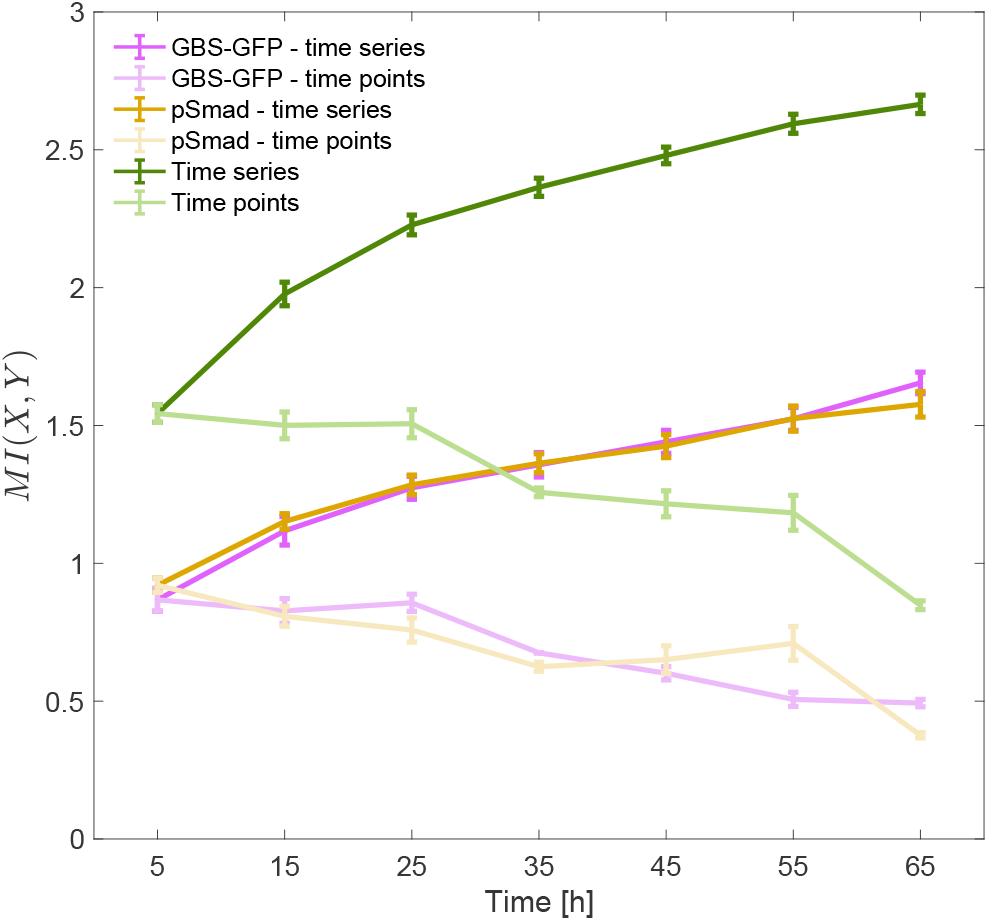
Mutual information (MI) between the 15 positions along the dorsal-ventral axis and the levels of the signals GBS-GFP and pSmad1/5 measured at these positions. The values of MI are first computed using the levels of GBS-GFP and pSmad1/5 separately, and then based on the levels of the two signals combined (where it is not otherwise denoted). The levels were measured at 7 time points during development. The time-point mutual information was computed using signal levels measured at each time point individually. The time-series mutual information is computed using the series of signal levels from the first time point up to each later point. The values are averaged over different embryos at a single time point and over different combinations of embryos when time series is considered. The error bars are the standard deviations. Units are bits.

### Temporal Information from Morphogen Gradients Increases Positional Information

The time-series mutual information increases with the inclusion of an increasing number of time points, indicating that new information continues to accumulate over time. The information accumulated during the first 65h from only Shh signaling is sufficient to distinguish 3 regions along the neural tube. For pSmad1/5 activity, the *MI* values are similar. By contrast, the values of mutual information for the combined Shh and BMP signaling indicate that the information accumulated during the first ∼30h is sufficient to resolve 5 regions without error (2^2.32^ ≈ 5). This rises to 2.67 *±* 0.03 bits by 65h (2^2.58^ ≈ 6). This confirms that the two morphogens acting together carry significantly more information. Overall, the analysis indicates that the combined signaling of the two morphogens over time changed the result from no two regions resolved at a single time point to almost 7 distinguishable regions accumulated over 65h.

Since positional information is used to classify cells into correct domains of gene expression, it is reasonable to ask what the precision is in terms of number of cells. Using the interpretation of 2^*MI*(*X,Y*)^, the time-series mutual information for the combined signaling revealed that the average absolute length of the regions that can be perfectly distinguished, in terms of average cell diameters, remains approximately the same during the first ∼30 hours, around 6 cell diameters. After that, the length increases; it is ∼7 cells for the 35h time course and ∼12 cells for the 65h of signaling. If we used the time-point mutual information, this length would be 6, 8, 10, 16, 20, 26, 42 at the seven time points, respectively. This means that during the first ∼30h of development, the average amount of information that is accumulated from Shh and BMP signaling increases proportionally to the tissue growth, maintaining positional precision. This result is in agreement with the claim that the pattern is established during the first ∼30h, which was supported by the analysis of Zagorski et al. (3) and the experimental evidence of Kicheva et al. (6).

This result is dependent on the choice of the time points and this level of precision is lost when we consider the points every 20h, 30h or 60h (SI Fig. 3). As we decrease the time step, the total amount of information for the 65h time course significantly increases. We expect that for a sufficiently small time step, reducing the step further will not add new information. However, we do not have data to determine the exact time step at which this happens. Here, we assume that the signal levels received over time are stored. This comes with a cost, which increases with the dimensionality of the time series, which is, on the other hand, limited by the memory capacity of the cells (35). Another disadvantage of longer time series is that, since the time points are closer, there will be more redundant information (35). In our case, considering all the 7 time points leads to a high level of redundant information (SI Fig. 4), which raises the question of the efficiency of encoding information in this way.

We expected the time-series information to increase with time, and we noticed that the increase in the information becomes smaller at each next time step (SI Fig. 5). The results show that the most information is gained early in the development, when the gain is also higher with a smaller time step. This is true for the time points 5h, 15h and 25h, or approximately the first 30h of the development. Later, as the redundancy increases, larger time steps bring higher information gain.

## Comparison of Positional Information from Time Series and Time Integrals

The dynamics of neural tube patterning challenges both the concept of positional information and the information-theoretic approach to studying dynamic signals. If the duration during which signal levels are maintained above a particular threshold determines position (26), the question is whether a whole vector of signaling dynamics should represent encoded positional information. It has been suggested that positional information in the neural tube could be encoded as the time integral of signaling (26, 28). Therefore, we considered cumulative level and duration by summing the entries of vectors for each of the two signals and computed *MI*(*X, Y*) where a single output response corresponding to an input *x*_*i*_ is now 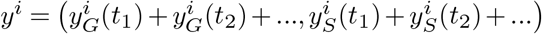.

This time-integral *MI* during the first ~ 30h is slightly lower than the time-series mutual information, Fig. 4. The difference between the two slowly increases over time since the time-integral mutual information plateaus after 35h, with even a slight decrease at 65h. If the domains of gene expression are established during the first∼30h, any new information that arrived after that time would not be used.

**Fig. 4.**
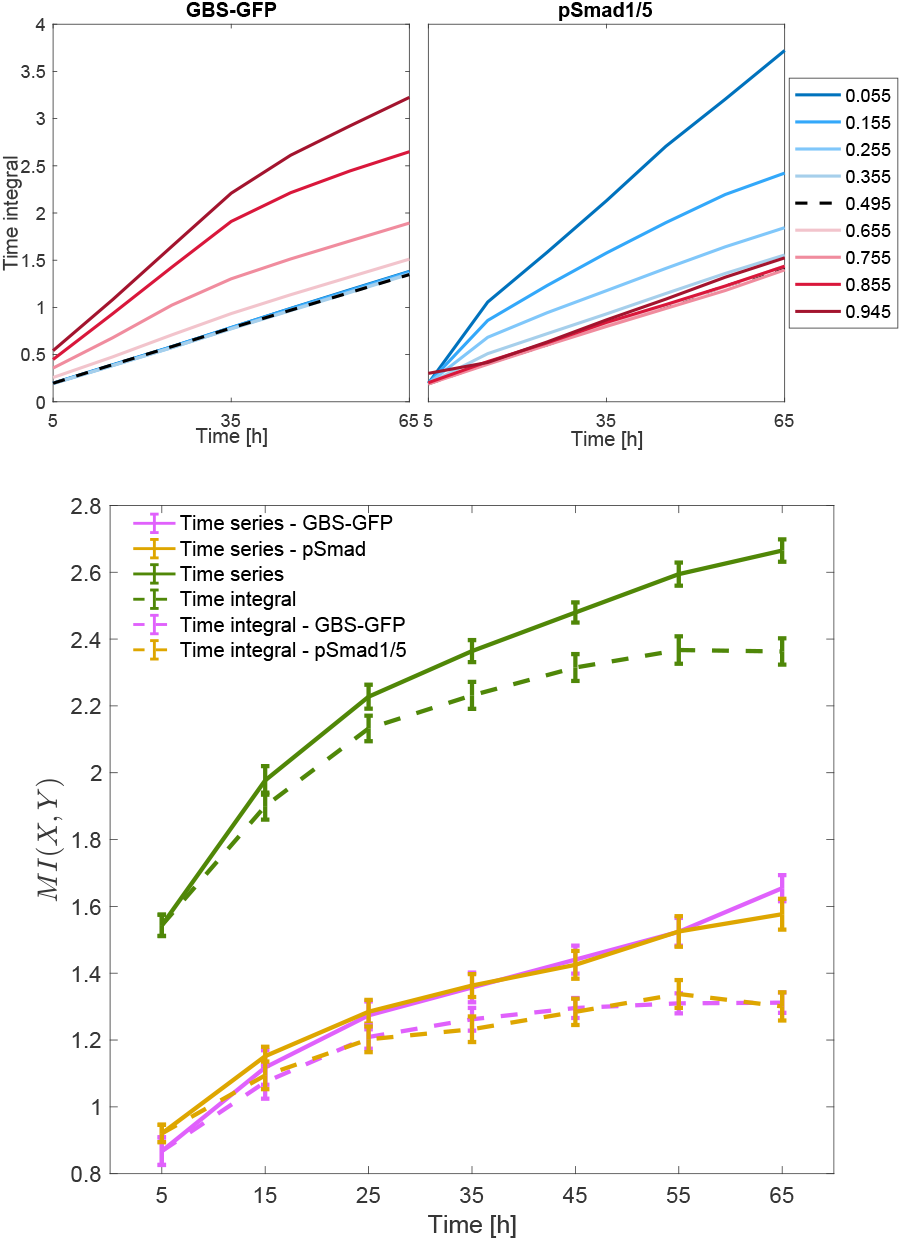
Top: the mean values of time integrals of GBS-GFP and pSmad1/5 over time at the indicated relative positions. Bottom: the time-series and the time-integral mutual information using the levels of GBS-GFP and pSmad1/5 separately, and based on the levels of the two signals combined. The time-integral mutual information is measured by summing the signal levels received over time, up to each time point. Units are bits.

Furthermore, when considering the two signals together, the time-integral mutual information gives approximately the same level of precision during the first∼30h as the time-series mutual information, i.e. the absolute length of the regions along the neural tube that can be perfectly distinguished is ∼6 cells during this time. Later, the length increases and is around 15 cells by 65h.

Encoding information using combined dynamics of multiple factors has also been considered by Granados et al. (34). They introduced a measure for the degree of redundancy between a pair of transcription factors which gives 0 when they are completely independent and 0.5 when they are completely redundant. In the time-series scenario, the redundancy between the activities of GBS-GFP and pSmad1/5 is approximately 0.13 during the first 45h and then increases to 0.18 by 65h. When considering time integrals of the signals, the redundancy decreases over time to less than 0.1 by 65h (SI Fig. 6).

These results support experimental findings (26) which suggested that the time integral is a suitable mechanism for encoding positional information in the neural tube.

Although we do not know precisely what features of morphogen signaling cells use to obtain knowledge about their positions, the introduction of time led to considerably more information than if static morphogen gradients were considered. This may explain why developmental systems have been found to respond to duration of morphogen signaling as well as its level. It emphasises the importance of a definition of positional information that captures this extended morphogen sensing.

### Probability Distributions of Positional Assignments

The number of bits does not necessary indicate the number of regions that are perfectly distinguishable in the neural tube: 1 bit could indicate that two regions are completely resolved or it could indicate that 3 or more regions are resolved, but with some error (38). As mentioned above, SLEMI tells us how well the true position of an observed signal level can be inferred by estimating the probabilities 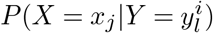, for all *i, j* = 1, …, 15 and *l* = 1, …, 102, where 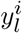 denotes *l*^*th*^ output response measured in bin *x*_*i*_. Here, a signal level 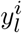 can be either a scalar, when the level of GBS-GFP or pSmad1/5 is taken separately at a single time point or its time integral, or a vector when both signals and/or consecutive time points are considered. Since signal levels are measured at 90 relative positions, every 0.01 of relative length from position 0.055 to 0.945, we find the average probability of each of these positions being in any of the 15 bins by averaging 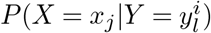 over signal levels measured at that position (21). We denote these probabilities by *P* (*X*^∗^ = *x*_*j*_ *x*^*i,k*^) where *X*^∗^ is a discrete random variable with values *x*_1_,…, *x*_15_ corresponding to the bins and *x*^*i,k*^, for *i* = 1, …, 15 and *k* = 1, …, 6, denotes one of 6 relative positions in a bin *x*_*i*_.

Using the levels of GBS-GFP and pSmad1/5 activity together at individual time points, the probabilities of classifying relative positions into correct bins along the patterning axis increase at the poles of the neural tube and decrease everywhere else with each next time point (Fig. 5, SI Fig. 7). The lack of scaling of the gradients as the neural tube grows leads to this expansion of the region of low positional information over time (Fig. 2). On the other hand, the time-series analysis shows that information accumulated during the first 35h leads to notably higher probabilities of correct positional assignment along the whole length of the neural tube, particularly for the positions within the dorsal and ventral third. Considering the time integral of signal levels, the information along the neural tube is distributed in the same way as in the time-series scenario. The time integrals of signal levels lead to slightly lower probabilities during the first 35h, compared to the time-series scenario, and this difference increases over time. More precisely, the highest difference between the time-series and time-integral probabilities after 15h, 25h and 35h is 0.06, 0.09 and 0.09, respectively, while after 65h it is 0.27 (SI Fig. 8).

**Fig. 5.**
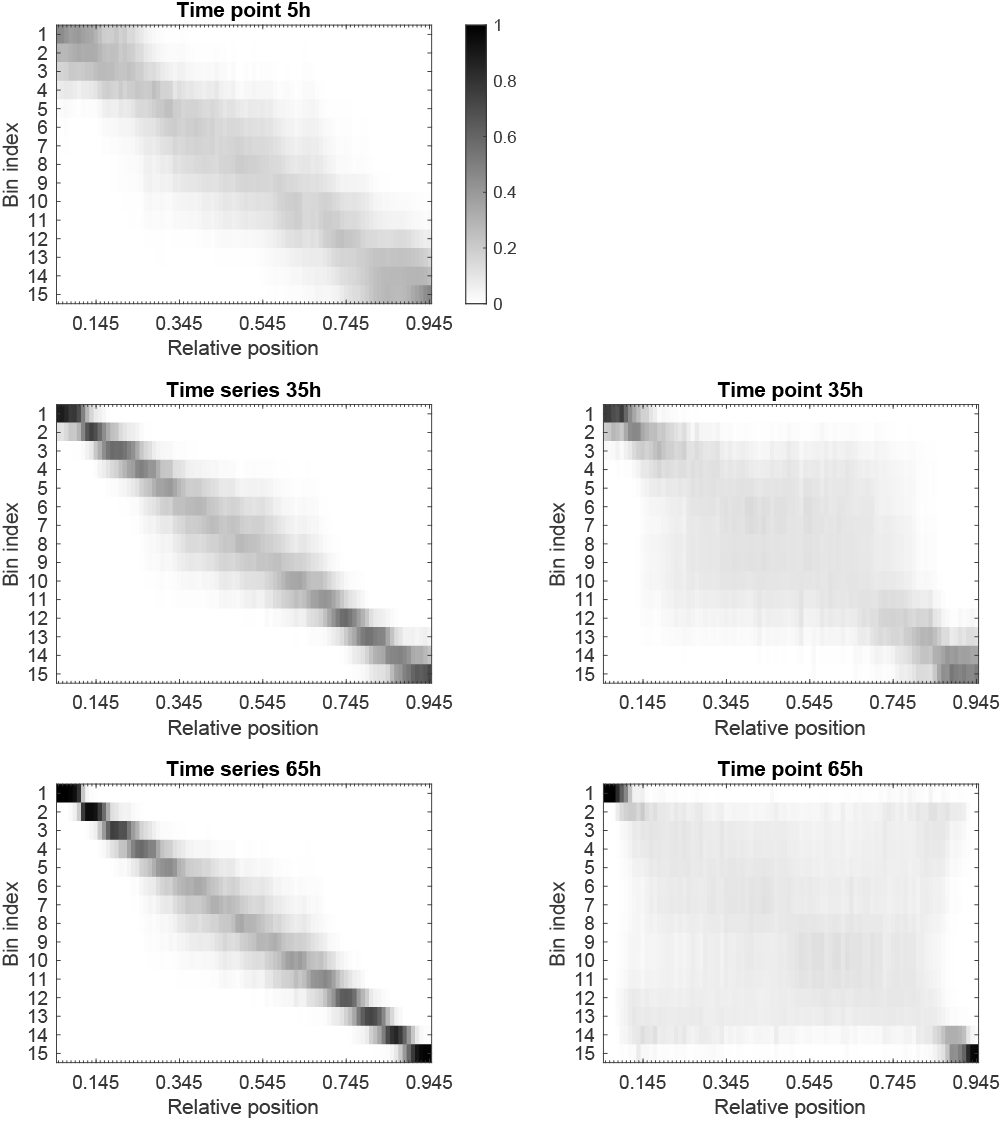
The average probability of being in any of the 15 bins after observing levels of the two signals measured at one of the relative positions from 0.055 to 0.945, at time points 5h, 35h and 65h, and for the time series of levels up to 35h and 65h.

We looked closely at the average distribution *P* (*X*^∗^|*x*^*i*^), the probabilities of being in any bin after observing a signal level measured at a single relative position *x*^*i*^ within bin *x*_*i*_, for *i* = 4, 8, 12 and *x*^4^ = 0.255, *x*^8^ = 0.495 and *x*^12^ = 0.755, at time point 5h and for time series and time integral over the first 25h, Fig. 6.

**Fig. 6.**
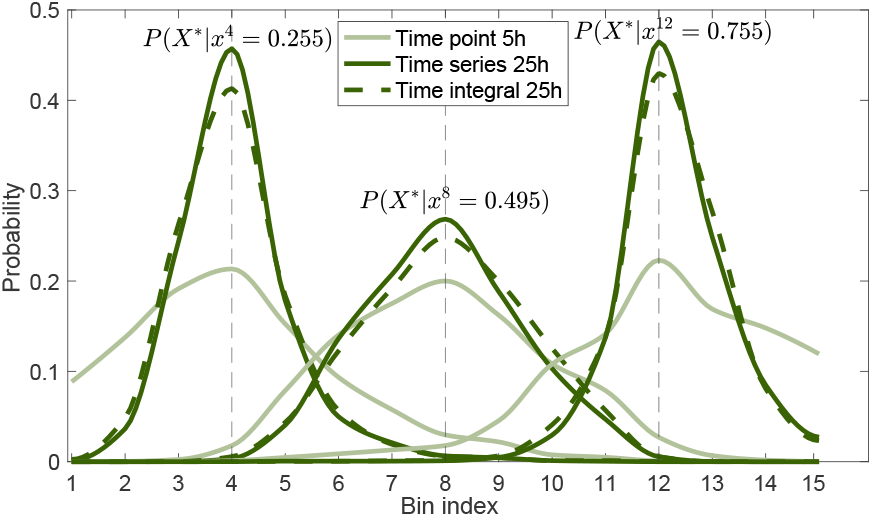
The probability distributions *P* (*X*^∗^ | *x*^*i*^) showing the probabilities of being in any bin after observing a signal level measured at a relative position *x*^*i*^, for *x*^4^ = 0.255, *x*^8^ = 0.495 and *x*^12^ = 0.755. We computed these at time point 5h and for time series and time integral over the first 25h.

The increase in the probability of being in the correct bin between 5h and 25h is particularly noticeable for the positions near each pole of the neural tube, where one signal is maintained above a certain threshold during the first 25h, while the levels of the other signal take the lowest values (Fig. 2). This is consistent with the findings that the duration of signaling influences cell fate decisions. For the middle of the neural tube, where both signals take the lowest values, information also accumulates over time and the probabilities of correct discrimination increase, although to a notably lesser extent resulting in the less marked increase in the peak of the distribution.

Observing the time-series probabilities for the 35h time course obtained from the levels of GBS-GFP and pSmad1/5 activity separately, it is evident that the signaling from the second morphogen contributes significantly to precision along the major part of the neural tube, Fig. 7. Shh signaling alone provides insufficient information for the region [0.055, 0.585] of the neural tube, and BMP signaling provides limited information to [0.445, 0.945], where the probabilities of correct discrimination are around 0.1.

**Fig. 7.**
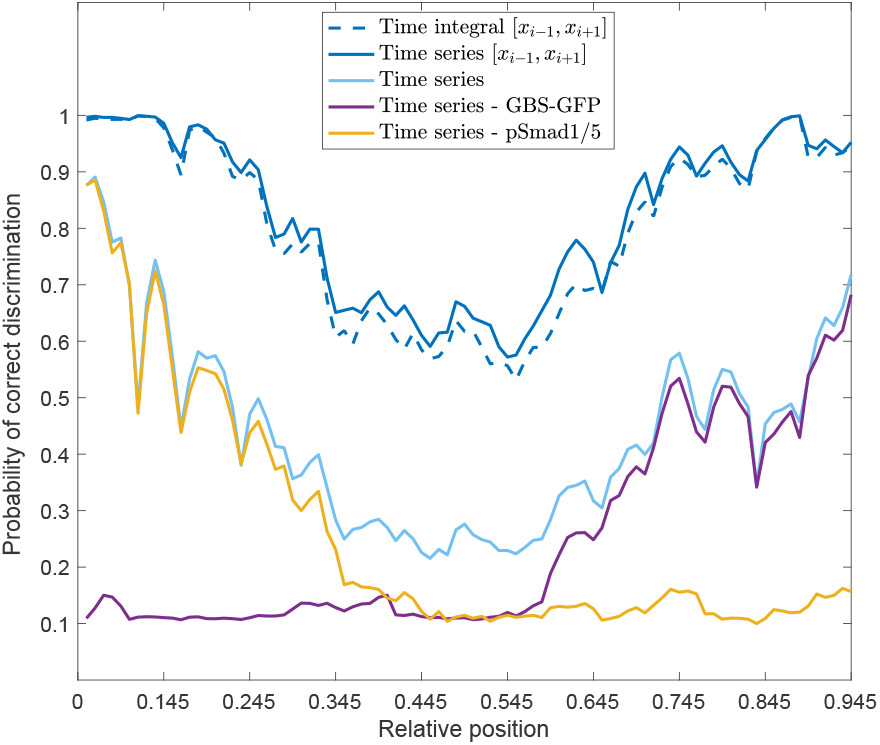
The average probabilities of each of the relative positions being correctly classified in its own bin (where it is not otherwise denoted) or in the region containing the correct bin and its adjacent bins, denoted by [*x*_*i*−1_, *x*_*i*+1_]. All probabilities are calculated on the basis of the levels of the signals during the first 35h, using the time series or the time integral. When the probabilities are obtained using the levels of the two signals separately, it is denoted which signal is used. Otherwise, they are obtained based on the two signals together.

### Spatial Patterns of Positional Uncertainty

A signal level 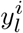 is correctly decoded if the probability distribution 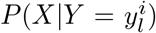 attains its maximum at the correct bin *x*_*i*_, and incorrectly if the maximum is at any other *x*_*j*_, *j* ≠ *i*. We assume that this maximum is always unique since the system should provide unambiguous positional information. Therefore, the fraction of incorrectly decoded signal levels at time points 5h, 15h and 25h is 32%, 30% and 34%, while for the 15h, 25h, 35h and 65h time courses it is 16%, 12%, 11% and 5%, respectively. When we consider the time integrals after the first 15h, 25h and 35h, this fraction is 20%, 17% and 19%. Examining the relative positions that are incorrectly decoded at the time point 5h indicated that these positions are almost uniformly distributed along the axis, with the most incorrectly discriminated positions at the ventral end, Fig. 8. Taking the time series and the time integral after the first 25h decreases the number of incorrectly decoded levels in the ventral and dorsal third of the neural tube, while in the middle only the time series results in a significant improvement.

**Fig. 8.**
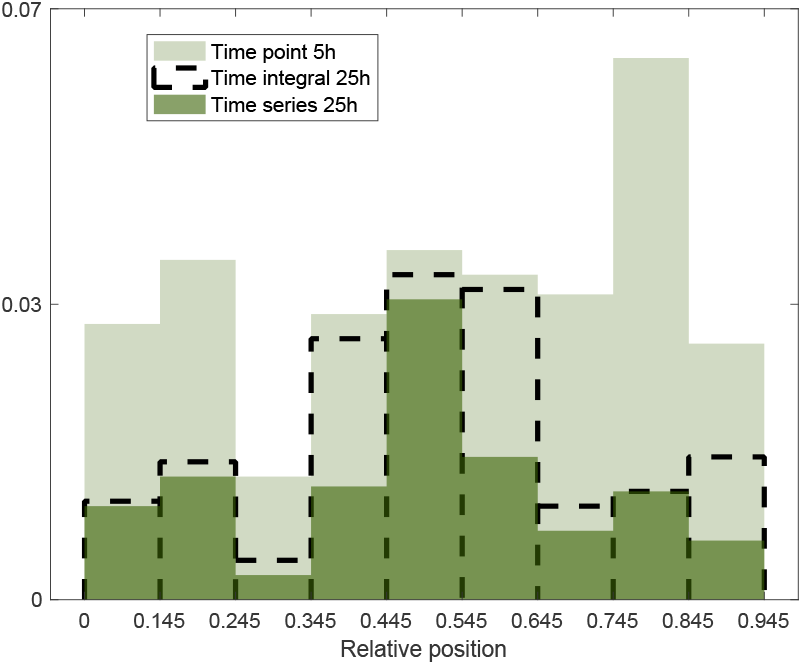
The distributions of incorrectly decoded signal levels obtained at time point 5h and using the time integral and the time series during the first 25h. The *x*-axis shows the relative positions where the incorrectly decoded signal levels are measured and *y*-axis the fraction they make out of all the signal levels considered at the given time. The results are obtained on the basis of the two signals together.

The time-series and time-integral results showed that after 25h and 35h a correct bin *x*_*i*_ is mistaken for a bin adjacent to it, i.e. for *x*_*i*_^−^_1_ and *x*_*i*+1_ (or only *x*_*i*+1_ if *i* = 1 and *x*_*i*_^−^_1_ if *i* = 15). In terms of relative length, this is within the region *±*0.06 from the correct bin, which is ∼2.5 cell diameters at 35h. On the other hand, at the first three time points individually, 10%, 15%, 21% of incorrectly decoded signal levels, respectively, are most probable to be in bins that are not adjacent to the correct one. We calculated the average probability of being inside an area including *x*_*i*−1_, *x*_*i*_ and *x*_*i*+1_ – the true bin and its nearest neighbours – after observing a signal level measured at each of the considered relative positions, found as the sum *P* (*X*^∗^ = *x*_*i*−1_|*x*^*i,k*^) + *P* (*X*^∗^ = *x*_*i*_|*x*^*i,k*^)+*P* (*X*^∗^ = *x*_*i*+1_|*x*^*i,k*^). We denote this sum as *P* (*X*^∗^∈ [*x*_*i*−1_, *x*_*i*+1_] | *x*^*i,k*^). Compared to the average probabilities of being in the correct bin, the average probabilities of being in the region [*x*_*i*−1_, *x*_*i*+1_] take considerably higher values. In particular, considering the time integral and the time series over the first 35h, the average probabilities of being in the region [*x*_*i*_ ^−^_1_, *x*_*i*+1_] are higher than 0.5 for all positions, and for the dorsal region [0.055, 0.255] and the ventral region [0.695, 0.945] these probabilities are higher than or equal to 0.9, Fig. 7. If we assume that *P* (*X*^∗^ | *X* = *x*^*i,k*^) is a normal ∼2 cell diameters during the first ∼30h, as its mean, we can approximate its standard deviation. At the first four time points individually, the highest errors are found in the middle of the neural tube and are equal to ∼2, 3, 4 and 7 cell diameters, respectively. At the same points, in both the time-series and time-integral scenarios, the errors are ∼2, 2, 2 and 3 cells, respectively. This is consistent with the idea that signaling dynamics improves precision and maintains it at approximately the same level during the first ∼ 30h.

## Discussion

The development of the neural tube requires precise spatial patterning along the dorsal-ventral axis. This is governed by Shh and BMP morphogen gradients that provide positional information via intracellular signaling pathways (25). Using experimentally measured levels (3) of the downstream transcriptional effectors of these signaling pathways – GBS-GFP and pSmad1/5 activity, respectively – we used the SLEMI algorithm (32) to quantify and decode the positional information transmitted by the two pathways. This analysis confirmed that a single morphogen gradient contains insufficient information to accurately resolve positions, except in the most dorsal or ventral regions of the neural tube, adjacent to the source of morphogen. By contrast, the combination of Shh and BMP signaling provides substantially more spatial information. Furthermore, we found that both the temporal trajectories and the time integrals of the signals carried more information than instantaneous snapshots. Although temporal trajectories carry the most information at all times, both methods of encoding information using temporal profiles provide very similar results during the first 35 hours. Later only time series contribute new information, although at the cost of high redundant information. However, in both cases, during the first 35 hours there is sufficient information for accurate positional estimates, particularly for the dorsal and ventral thirds of the neural tube. Thus, cells would gain significantly more information by tracking signaling over the initial 35 hours of neural tube development and this would allow a substantially increased precision of positional decoding in the neural tube. Overall, therefore, tracking the temporal profile of the two signals improved precision, and the analysis is consistent with the temporal profile of the two signals together providing more positional information than either the signal individually or snapshots of the signals.

Nevertheless, in the middle third of the neural tube, where both BMP and Shh signaling is low, the information provided by the measured signaling gradients remained limited. While our analysis focused on Shh and BMP signaling, it is important to note that other morphogens may contribute to neural tube patterning. Retinoic Acid (RA) signaling from the paraxial mesoderm has been implicated in neural tube patterning, particularly in the intermediate regions and could provide additional positional information (39). Moreover, the Wnt family of secreted glycoproteins have been suggested to play a role in dorsal neural tube patterning and could provide additional information in conjunction with BMP signaling (40). The contribution of these additional morphogens to positional information, especially in the middle regions of the neural tube, warrants further investigation and integrating data on these signaling pathways into our information-theoretic framework could provide additional insight into neural tube patterning.

Previously Zagorski et al. (3) established a mathematical framework for decoding positional information in the neural tube. This involved several approximations. The probability distribution of signal at a single relative position *x, P* (*Y* |*X* = *x*), was assumed to be Gaussian. The signals were also assumed to be independent and only individual time points considered. Our approach relaxes these assumptions. Nevertheless, the conclusions are broadly consistent with the previous analysis. Similar to the results reported here, Zagorski et al. show that the time-point positional error is highest for the middle of the neural tube and increases over time; they found the highest positional imprecision to be ∼3, 4, 5, 8 cells at 5h, 15h, 25h, 35h, respectively. Moreover, Zagorski et al. conclude that a combination of signaling provides substantially lower positional error throughout the neural tube than either signal individually. The analyses are consistent with previous experimental evidence that the initial 30h of mouse neural tube development are critical for tissue patterning. Our approach extends the analysis from Zagorski et al. by taking series and integrals of signal levels during the initial ∼30h to conclude that the signaling over time decreases positional error. Patterning specifies 11 domains of neural progenitors. If these domains were of the same size, the length of each would be ∼3 cell diameters at ∼30h. We found that, at this point in patterning, the accumulated information is sufficient to perfectly distinguish regions containing ∼6 cells on average. Moreover, the highest error, given by the standard deviation, with which a bin of size less than ∼2 cell diameters can be specified is ~ 2 cells. Therefore, a domain containing 3 cells could be specified with an error of less than 2 cells. Zagorski et al. have measured the boundaries of the two domains of gene expression, Pax6 and Nkx6.1, which are positioned in the middle of the tube. The precision of the boundaries was found to be approximately *±*1 and *±*2 cells at 30h. As suggested by Zagorski et al. (3), the similarity in precision of signaling profiles and boundary positions indicates that cells decode the information in a way that minimises imprecision.

Our results show the discrepancy in precision between the middle and the ends of the neural tube, which raises the question if this is potentially compensated for by another source of information or another mechanism for information decoding. The gene regulatory networks downstream of morphogen signaling play a crucial role in the interpretation of positional information (2, 25). It is still unknown how precisely the levels of Shh and BMP signaling are measured downstream. If only thresholds of their concentrations can be sensed (41), the question is how much positional information is used. However, cross-repressive interactions between transcription factors help refine boundaries, as evidenced by shifting, noisy boundaries when specific transcription factors in the downstream network are lost (5). The design of the network appears to enhance patterning precision by reducing the effects of transient fluctuations in signaling. Extending the information-theoretic approach directly to quantified transcription factor levels could provide further insight into how the gene regulatory network processes time-varying positional information and affects the precision of tissue patterning. In addition, corrective mechanisms such as differential adhesion have been implicated in contributing to patterning precision (24). Assessing the role such mechanisms play in the precision of neural tube patterning would provide insight into the relative contribution of the information provided by morphogen signaling compared to downstream mechanisms.

The information-theoretic approach provides a powerful framework for analysing morphogen-based patterning that does not rely on detailed mechanistic knowledge of the patterning process. The data-driven nature, however, means that the accuracy of the experimental measurements of signaling dynamics influences the analysis, and the analysis is constrained to the available time and spatial points at which measurements are made. In regions of the tissue where the level of signaling is low, the fluorescent intensity of the corresponding signal reporters is low and approaches background levels; this increases the uncertainty in estimated signaling levels and results in lower estimates of positional information than would be obtained if more sensitive reporters were available. In addition the temporal trajectories used in our measurements contain variations at the level of the whole population. This means that the computed bits of information do not represent the information provided to an individual single cell, but rather a general profile of positional information available in the system that can be used for patterning. It would be ideal to follow the level of signaling in individual cells in the same embryo over time. This would provide additional insight into how much information cells receive and allow quantification of the noise that arises in a single embryo. Moreover, analysing data from experiments that modulate different aspects of the Shh and BMP signaling pathways could provide a deeper understanding of the information transmission in the system. For example, in Zagorski et al. (3), signal gradients are measured in embryos that are hypomorphic for Shh signaling.

How to define position in a growing domain is still an open question (19). Following a single cell and its movement and division would give us the most accurate picture of the cell’s position over time. The growth in the neural tube has been studied (6, 42). However, how growth is coupled with morphogen signaling and gene regulatory networks remains to be clarified.

Overall, the study supports the conclusion that information theory combined with quantitative dynamic data is a powerful approach to unravel the mechanisms of precise spatial patterning in development, and permits a quantifiable definition of *positional information*. The temporal integration of morphogen signals is likely crucial for the patterning of many developmental systems, as the duration of morphogen signaling has been implicated in the patterning of other tissues (43–45). The methodology described in this study could therefore provide insight into other developmental systems. The success will depend on obtaining the appropriate quantitative data and expanding theoretical and experimental knowledge of the systems.

## ACKNOWLEDGEMENTS

We are grateful to Anna Kicheva and Marcin Zagórski for helpful comments and advice. AM’s work was supported by a teaching assistantship from the Department of Mathematics, UCL. Work in JB’s group is supported by the Francis Crick Institute, which receives its core funding from Cancer Research UK (CC001051), the UK Medical Research Council (CC001051), and the Wellcome Trust (CC001051); and by the Wellcome Trust (220379/D/20/Z).

## Notes

### Competing Interest Statement

The authors have declared no competing interest.

### Summary of Updates

Some clarifications throughout the main text, but mainly the Supplemental Information updated with results obtained using two new computational tools and their comparison with the one used in the main paper.

