## Supplementary Information for "Dynamics of positional information in the vertebrate neural tube"

**Data.** The data used in this work are from previously published work (1), they are not publicly available and were provided to us by the authors. Shh signaling activity was measured in the mouse neural tube using a transcriptional reporter, GBS-GFP, for the transcription effector of Shh signaling, Gli. BMP signaling activity was measured using an antibody specific for phosphorylated SMAD1/5, pSmad1/5, the activated version of the BMP transcriptional effectors. The embryo staging and analysis adopted by (1) was used: embryos were classified according to the number of somites, where one somite is generated every 2 hours in the mouse. The dorsoventral lengths of the neural tubes were fitted to $L = L_0 \exp(t/\tau)$ , where $t$ is the number of somites multiplied by 2h. The embryos were then reclassified based on the fit. Seven independent time courses were considered separately: the intensity profiles of GBS-GFP and pSmad1/5 in each time course were recorded and processed together to minimise variability (2). This resulted in measurements of GBS-GFP and pSmad1/5 at 7 time points, 5 hours, 15h, 25h, 35h, 45h, 55h and 65h. GBS-GFP and pSmad1/5 activity was measured along the dorsal-ventral length of the neural tube, every $0.01$ of relative length, starting from the relative distance $0.055$ at the dorsal end, where the first $5.5\%$ of the neural tube represented the source of BMP, to $0.945$, with the remaining $5.5\%$ considered the source of Shh. Further details can be found in (1).

To ensure a large enough sample of output measurements for each input value as required for SLEMI ( $n > 100$) (3), we divided the neural tube into 15 positional bins along the dorsal-ventral axis, and the corresponding output values are levels of GBS-GFP and pSmad1/5 measured within these bins, Fig.1. As was mentioned, the signals are measured in an embryo only once and the number of embryos considered at each time point is different, Fig. 9. The number is smallest for the time point 35h, 17 embryos, which is the reason for taking bins containing 6 relative positions, which then results in $17 * 6 = 102$ different output measurements in a single bin for that time point, across 17 different embryos. For all other time points, we have more embryos, so we choose 17 different ones. We take 20 different combinations of 17 embryos for these time points and compute mutual information in each case. We plotted the average values of mutual information, and the error bars are the standard deviations.

To find estimation bias due to the finite sample size, we use the method of shuffle control where we randomly permute the bin indices of the measured signal levels to check if the value of mutual information will be zero (4). The values we obtain in the time-series scenario for the two signals together increase from $0.01$ to $\sim 0.1$ after 65h, Fig.10. This value should be subtracted from the obtained values of mutual information, but they do not significantly affect the results.

**SLEMI.** To compute mutual information, we used a relatively new framework proposed by Jetka et al. (3), SLEMI - statistical learning estimation of mutual information. SLEMI substantially simplifies the calculation of $MI(X,Y)$ for the time-series output and avoids the estimation of high-dimensional probability densities $P(Y|X=x_i)$. The first step is expressing $MI(X,Y)$ in the following form

$$MI(X,Y) = \sum_{i=1}^{m} P(x_i) \int_{\mathcal{Y}} P(y|X=x_i) \log_2 \frac{P(x_i|Y=y)}{P(x_i)} dy.$$

The integral in the above formula is equivalent to the expected value of the term $\log_2 \frac{P(x_i|Y=y)}{P(x_i)}$ with respect to the probability distribution $P(Y|X=x_i)$. Since the data containing all output measurements for the input value $x_i$ is a sample from this distribution, for a large enough number of measurements ($n$), we can use the Law of Large Numbers (LNN) and approximate the integral by the average with respect to the data. In that way, the form of $MI(X,Y)$ that SLEMI computes is obtained:

$$MI(X,Y) = \frac{1}{n} \sum_{i=1}^{m} P(x_i) \sum_{l=1}^{n} \log_2 \frac{P(x_i|Y=y_l^i)}{P(x_i)}$$

We know that $P(X)$ is uniform, so, now, we are missing only $P(x_i|Y=y_l^i)$ to compute $MI(X,Y)$, and Jetka et al.(3) propose to approximate $P(x_i|Y=y)$ using *logistic regression*. The model is based on the assumption that log of the ratio of the probability of observing $y$ belonging to class $i$ to the probability of observing $y$ belonging to class $m$ is linear with respect to $y$:

$$\log\left(\frac{P(x_i|Y=y)}{P(x_m|Y=y)}\right) \approx \alpha_i + \beta_i^T y \quad \textbf{(1)}$$

for all $i = 1,2,...,m-1$. The provided R package finds the values of parameters $\alpha_i$ and $\beta_i$ that best fit the data and derives each $P(x_i|Y=y)$. Further details can be found in (3).

DRAFT

Computing $MI$ when the output is given by temporal trajectories remains a challenging task. There are only a few computational tools that are able to do this using a given dataset. The most recognised ones are the method proposed in Selimkhanov et al. (5), which uses $k$-nearest neighbor ($k$NN) density estimator, and the decoding-based approach developed by Granados et al. (4) using a machine learning classifier called a Support Vector Machine (SVM).

As explained above, the tool that we chose for our analysis, SLEMI, relies on the assumption given by the equation (1). In order to validate our results and check that this method is suitable for our data, we computed $MI$ using the other two tools, the $k$NN-based and decoding-based approaches, which do not depend on the same assumption. The values of $MI$ computed using all three tools are presented in Fig.2. The tools give similar results and do not change the conclusions of our main paper. The $k$NN method gives almost the same values of $MI$ as SLEMI when the two signals, GBS-GFP and pSMAD1/5, are considered separately. This comparison is done only in the time-series case, since kNN can only be applied to vectors. On the other hand, for each signal, the decoding-based method leads to higher values of $MI$ compared to SLEMI, although this difference does not exceed $0.2$ bits. The same is noticed in the time-point scenario for the two signals combined. However, when we consider the time series of the two signals together, both $k$NN and decoding-based method give lower values than SLEMI after 35h, with the lowest values coming from the decoding-based one. It has been argued that $k$NN and decoding-based method might not distinguish different dynamical patterns as well as SLEMI (6), which could be the reason behind this result. For the time integrals of the two signals, all three tools give very similar values of $MI$.

Jetka et al. (3) demonstrated the advantages of their SLEMI algorithm over the $k$NN-based approach. They created synthetic datasets with available exact solutions and concluded that SLEMI performed better then $k$NN. This was especially true for smaller sample sizes and higher dimensionality of the output, which describes the data that we are analysing. The results also depend on the right choice of the parameter $k$, which is usually difficult to achieve. We decided to use $k = 10$, as was done in both Selimkhanov et al. (5) and Jetka et al. (3). The kNN method may also be inaccurate for vectors whose dimension is higher than 10 (6), which would be the case for the 55h and 65h time courses of the two signals combined.

The decoding-based approach and SLEMI both treat approximation of probabilities required to compute $MI$ as a classification problem in machine learning. However, SLEMI estimates the probability of an output response being in any of the input classes, $P(x_i|Y = y)$, while the decoding-based approach computes the probability of correct or incorrect classification in terms of the distribution $P(\hat{X}|X = x_i)$ , where $\hat{X}$ represents input values decoded from the output responses. Hence, the decoding-based approach estimates $MI(\hat{X}, X)$ which represents a lower bound to the exact information $MI(X, Y)$ (7). Good performance of this tool has not been confirmed when the number of different input values is greater than 5. Since this number in our analysis is 15, we are not aware what kind of bias this can potentially introduce to our results (6).

The data is prepared in the same way for each tool, as described in the previous section. The computations are performed 20 times using different combinations of embryos, which resulted in the mean values plotted in Fig.2 with error bars representing standard deviations. The results of the decoding-based method also include the mean and standard deviation of the estimator obtained using a bootstrapping procedure. This involves resampling the fraction of the dataset used to train the classifier (4).

The scripts used for $k$NN method are part of the SLEMI package (3), while the path to the decoding-based software can be found in (4).

Comparing the three tools, we decided to use SLEMI as the one with the least number of limitations that can be used for a broader range of datasets (6).

DRAFT

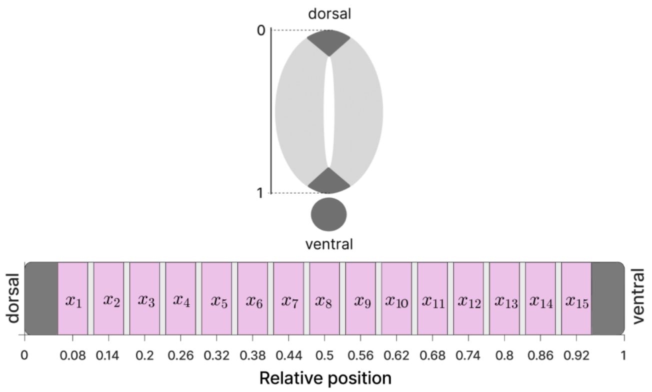


**Fig. 1.** The $15$ bins, of relative length $0.05$ each, along the relative dorsal-ventral length of the neural tube, from the relative position $0.055$ up to the position $0.945$, where the position $0$ corresponds to the dorsal end and $1$ to the ventral end. Below each bin, we show the relative position corresponding to the middle point of the bin.

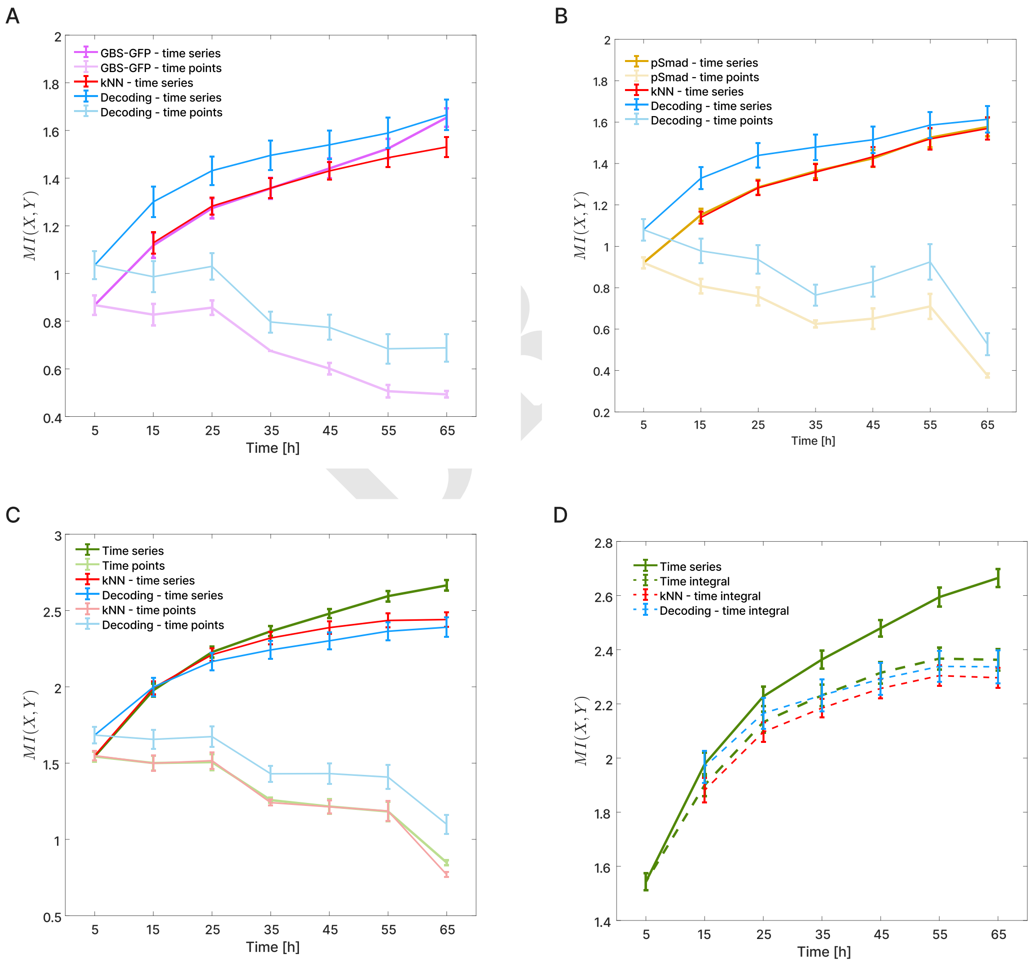


**Fig. 2.** Comparisons of the three tools for computing the mutual information, SLEMI, kNN and decoding-based methods, for: A) GBS-GFP in time-series and time-point scenario; B) pSmad1/5 signal in time-series and time-point scenario; C) the two signals combined in time-series and time-point case; D) the time integrals for the two signals together (compared with the SLEMI time-series values).

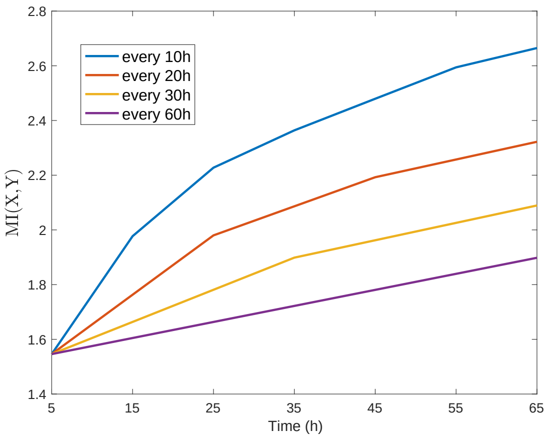


**Fig. 3.** The time-series mutual information when taking time points with different sizes of time steps , every $10$h (considered in the main paper), every $20$h, every $30$h and $60$h.

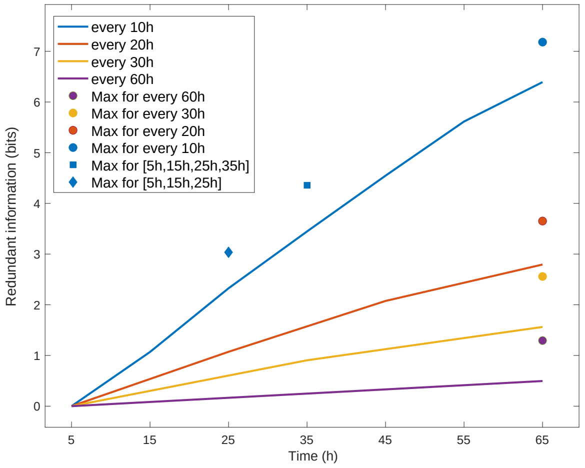


**Fig. 4.** The redundant information in the time-series mutual information, considering different sizes of time steps, every $10$h (used in the main paper), every $20$h, every $30$h and $60$h. It is computed using the method from Potter et al. (8) where they define the redundant information as the difference between the sum of the values of the time-point mutual information at the considered time points and the mutual information from the time series of those points. They also defined the upper bound for the amount of redundant information by $\frac{(k-1)}{k}\sum_{i=1}^{k} MI_{t_i}(X,Y)$, where $k$ is the number of considered time points and $MI_{t_i}(X,Y), i=1,\ldots,k$, the time-point mutual information at those points.

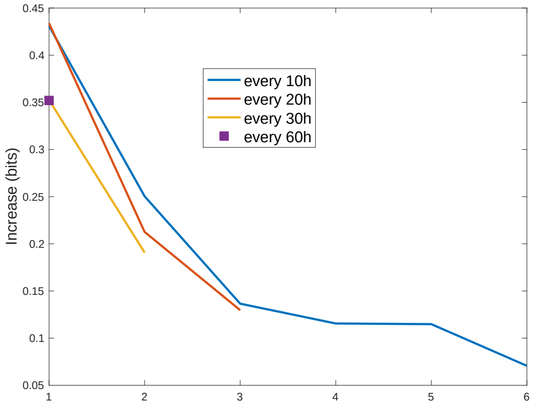


**Fig. 5.** The increase in the time-series mutual information at each next time point, considering different sizes of time steps, every $10$h (used in the main paper), every $20$h, every $30$h and $60$h. The $y$-axis indicates the order of the time step at which the increase is computed in a time series.

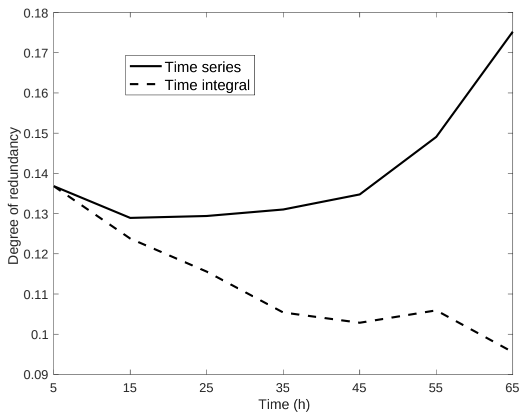


**Fig. 6.** The degree of redundancy between the two signals, GBS-GFP and pSmad1/5, in the time-series and time-integral mutual information, up to each time point. It is measured using the following formula from (4):
$1-\frac{MI(X,Y=\{G,S\})}{MI(X,Y=G)+MI(X,Y=S)}$, where $G$ and $S$ denote GBS-GFP and pSmad1/5, respectively, and $Y=\{G,S\}$ and $Y=G$ or $Y=S$ the cases when the output is the levels of both signals together and when it is the levels of only one of the signals.

DRAFT

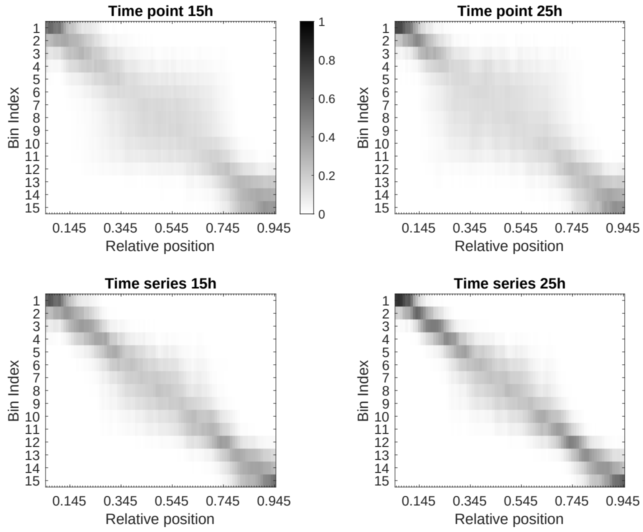


**Fig. 7.** The average probability of being in any of the $15$ bins after observing a signal level measured at one of the relative positions from $0.055$ to $0.945$. Probabilities are computed based on the levels of the signals measured at time points $15$h and $25$h separately, and based on the series of levels up to $15$h and $25$h . The probabilities are obtained based on the two signals together.

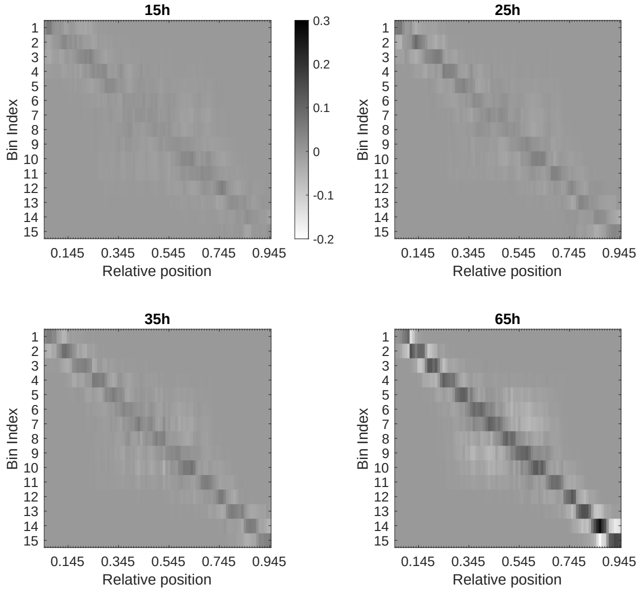


**Fig. 8.** The difference in the average probability of being in any of the $15$ bins after observing a signal level measured at one of the relative positions between the time series and the time integral scenario. Probabilities are computed based on the levels of the signals measured during the first $15$h, $25$h, $35$h and $65$h and the differences are found by subtracting the time-integral probabilities from the time-series probabilities. The results are obtained based on the two signals together.

| | GBS-GFP | pSmad1/5 |
|---|---|---|
| 5h | 30 | 24 |
| 15h | 62 | 40 |
| 25h | 69 | 46 |
| 35h | 17 | 19 |
| 45h | 42 | 55 |
| 55h | 32 | 33 |
| 65h | 21 | 19 |

**Fig. 9.** The number of embryos for each time point, for each of the two signals.

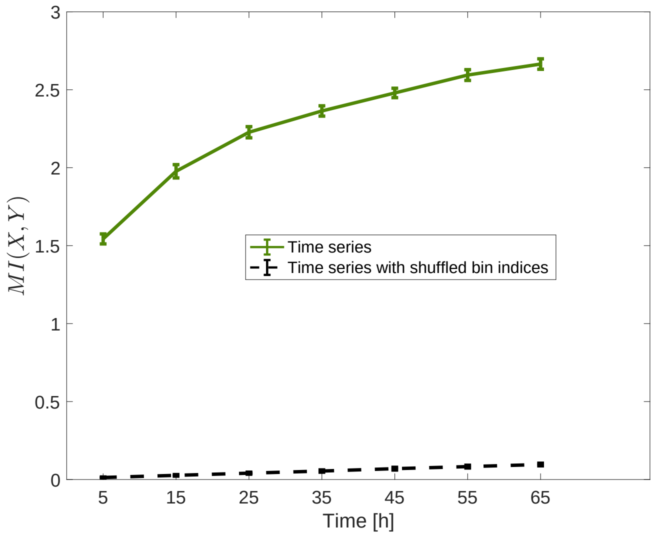


**Fig. 10.** The time-series mutual information with shuffled bin indices represents estimation bias due to finite sample size that should be subtracted from the original values of the time-series mutual information.


DRAFT
